# Individual dietary amino acid restrictions induce distinct metabolic and chromatin states

**DOI:** 10.1101/2023.12.06.570456

**Authors:** Spencer A. Haws, Yang Liu, Cara L. Green, Krittisak Chaiyakul, Pragyan Mishra, Reji Babygirija, Eric A. Armstrong, Anusha T. Mehendale, Irene M. Ong, Dudley W. Lamming, John M. Denu

## Abstract

Dietary protein and essential amino acid (EAA) restriction promote favorable metabolic reprogramming, although the extent to which shared or EAA-specific mechanisms facilitate diet-associated phenotypes remains unclear. Here, we compared the physiological and molecular effects of dietary methionine, leucine, or isoleucine depletion (Met-D, Leu-D, and Ile-D) in C57BL/6J mice. Each diet elicited responses not phenocopied by mTORC1 inhibition, including reduced fat mass and hepatic amino acid catabolism. Ile-D yielded additional distinct responses, highlighted by histone H2A/H4 hypoacetylation and maintained hepatic acetyl-CoA levels despite downregulated FA β-oxidation. Multi-Omics Factor Analysis of 14,139 data points objectively affirmed Ile-D phenotypes are distinct from Met-D or Leu-D and identified several metabolic and chromatin features as primary discriminators. Metabolic and epigenetic responses to Ile-D were recapitulated *in vitro*, suggesting underlying mechanisms represent fundamental cellular properties. Together, these results demonstrate EAAs can stimulate unique phenotypes and highlight distinct molecular mechanisms by which EAAs may inform metabolic health.

## Introduction

Dietary protein is a key regulator of metabolism, health, and longevity (^1^). It has long been suspected that the health benefits of a low protein diet arise not simply due to the consumption of reduced protein, but due to decreased levels of specific amino acids. For many years, attention has focused on the potential role of dietary methionine in regulating metabolism and health. Plant-based protein may have reduced levels of methionine compared to animal proteins, and vegans have lower than normal plasma levels of methionine; thus, many have attributed the health benefits of a vegan diet to decreased consumption of methionine (^2–4)^. The potential metabolic benefits of reduced methionine intake are readily observed in rodents, where restriction of methionine extends lifespan, promotes leanness and insulin sensitivity, and promotes the rapid loss of adipose tissue and improved glycemic control in diet-induced obese mice (^5–9)^.

However, methionine is only one of the twenty common amino acids, and there is a growing awareness that other dietary essential amino acids have critical roles in regulating metabolism and health. The most well-known of these are the branched-chain amino acids (BCAAs; leucine, isoleucine, and valine), which are elevated in individuals with diabetes, are associated with insulin resistance in humans and rodents, and which promote insulin resistance in rats when supplemented in the diet (^10^). Conversely, dietary restriction of BCAAs promotes glucose tolerance, insulin sensitivity, and reduces adiposity in mice and rats (^11–13)^. BCAA intake also regulates the lifespan of mice, with BCAA restriction extending the lifespan of mice and promoting lifelong leanness, and BCAA supplementation shortening lifespan and promoting obesity (^14,15^). Many of the metabolic and anti-aging effects of BCAAs may be mediated by isoleucine, as restriction of isoleucine is sufficient to improve metabolic health, extend lifespan, and rejuvenate aged mouse tissues (^16–19)^.

Fluctuations in dietary nutrient consumption have been proposed to regulate physiological responses, in part, via epigenetic mechanisms as central metabolites act as co-substrates for chromatin-modifying enzymes (^20,21^). For example, methionine is essential for the production of S-adenosylmethionine (SAM), which acts as a methyl-donor for methyltransferases. Reduced SAM availability is associated with both global and site-specific decreases in histone methylation post-translational modifications that play critical roles in regulating gene expression and genome stability (^22^). Similarly, BCAAs are catabolized to either acetyl-CoA (leucine), propionyl-CoA (valine), or both (isoleucine), and while having many other biological roles these BCAA-derived catabolites can also be used for the posttranslational modification of histones. Acetyl-CoA is required for histone acetylation, and acetyl-CoA derived from leucine can be utilized by the p300 histone acetyltransferase to acetylate multiple substrates (^23,24^). Isoleucine-derived propionyl-CoA can also be used as a substrate by histone acyltransferases and is required for propionylation of cardiac H3K23 (^25^). The effects of leucine and isoleucine are particularly interesting as recent work has shown that restriction of leucine and isoleucine have dramatically different effects on the metabolic health of mice, with restriction of isoleucine being necessary and sufficient for the metabolic benefits of protein restriction including reduced adipose tissue, increased energy expenditure, and improved glycemic control, while reducing leucine did not (^19^). Further, dietary levels of isoleucine, but not leucine, are associated with body mass index in humans (^19^).

It is thus clear that adaptive mechanisms exist which allow animals to actively respond not just to times of famine, but to decreased availability of specific essential amino acids. These may occur over different time periods, including meal to meal variation or simply during the day in response to feeding and circadian rhythm. Some of these mechanisms could involve the mechanistic Target of Rapamycin Complex 1 (mTORC1), an evolutionarily conserved sensor of amino acids - including Met, Leu, and Ile - that promotes growth and anabolic metabolism when amino acids are sufficient (^26^).

Here, we investigate the metabolic effects of restricting methionine, leucine, and isoleucine on the physiology and the liver metabolome, epigenome, and transcriptome of C57BL/6J male mice. We limited the treatments to 3 weeks, with the goal of capturing early-stage responses to depletion of EAAs. We further examine the potential role of mTORC1 in facilitating these responses by including a treatment group receiving an inhibitor of mTORC1, rapamycin. By assaying these treatment groups in parallel within a single study, we aim to quantitatively determine the degree to which dietary EAA restrictions elicit unique and/or shared physiological and hepatic responses to inform how the specific amino acid profile of dietary proteins plays a key role in regulating metabolic health at the molecular level.

## Results

### Essential amino acid restrictions elicit systemic physiologic responses unique from rapamycin treatment

To investigate how complete dietary restriction of select EAAs subsequently impacts different physiological measurements, we placed 14-week-old C57BL/6J male mice on one of four AA-defined diets for 3 weeks (**Fig. 1A**). Briefly, we utilized a widely used commercially available AA-defined Control diet in which 15.6% of calories are derived from AAs. We then generated a series of diets in which either methionine (Met-D), leucine (Leu-D), or isoleucine (Ile-D) were depleted. All of the diets are isocaloric, with identical levels of fat and carbohydrate; in each restricted diet, non-EAAs were increased to keep the calories derived from AAs constant (**Table S1)**. In parallel, a subset of Control-fed mice was treated daily with rapamycin in order to assess if mTORC1 inhibition mimics some or all of the physiological and molecular effects of EAA restriction.

**Figure 1:**
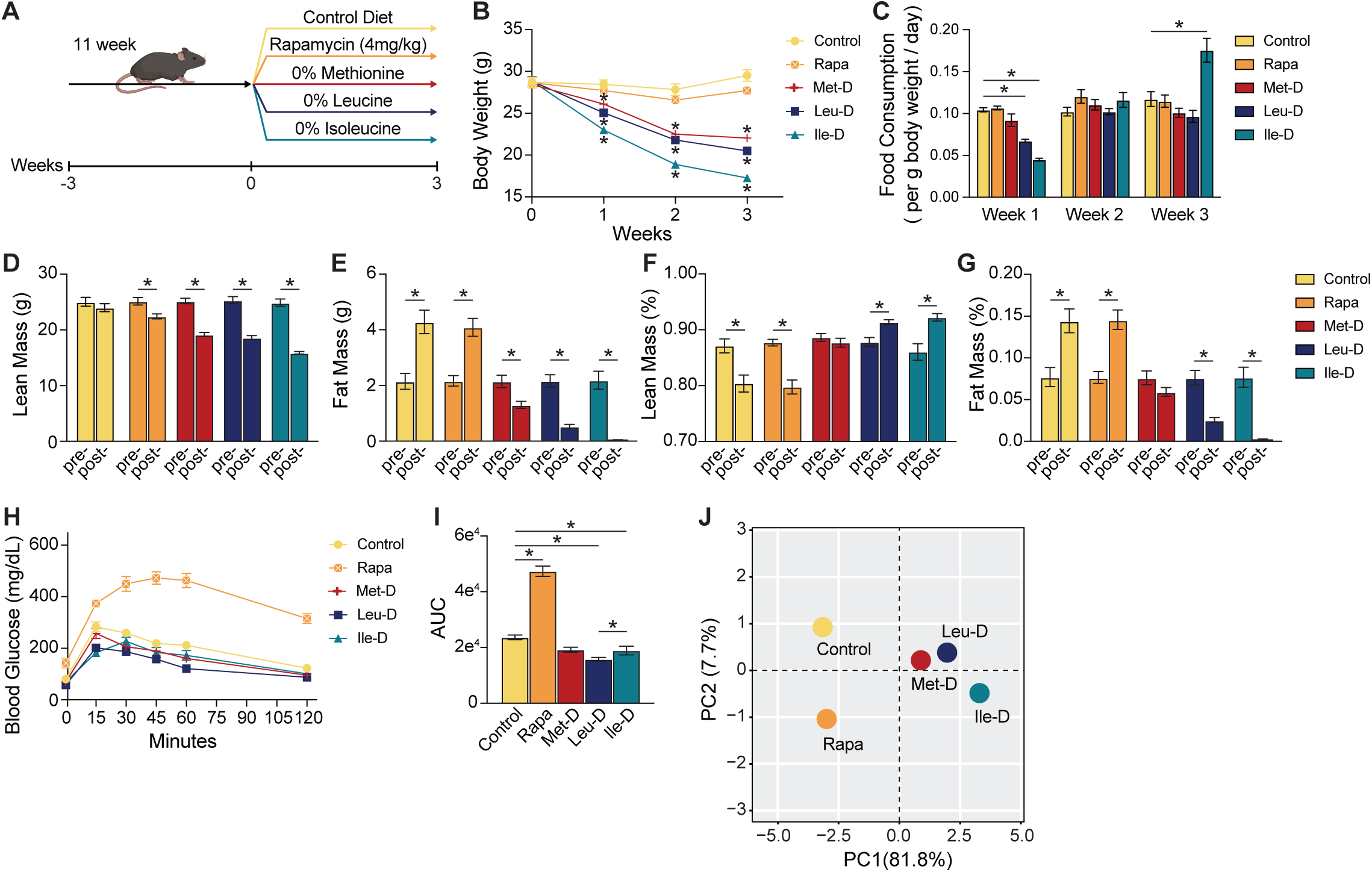
Physiologic responses to EAA restrictions and rapamycin treatment. (*A*) Visual representation of study design. (*B*) Weekly body weight measurements. (*C*) Bar graph illustrating weekly food consumption normalized to mouse body weight values. (*D-E*) Bar graphs depicting absolute changes in lean and fat mass measured via whole-body MRI following three weeks of dietary intervention or rapamycin treatment. (*F-G*) Bar graphs depicting the body composition percentages of lean and fat following three weeks of dietary intervention or rapamycin treatment. (*H-I*) Glucose tolerance test curves and calculated area under the curve (AUC) values. Error bars = SEM; N = 12; * = p-value < 0.05 as measured via Student’s t-test.

We found that restriction of EAAs, but not rapamycin treatment, led to a steady decrease in total body weight (**Fig. 1B**). Met-D and Leu-D mice exhibited a similar pattern of lost body weight, weighing 7.5 g and 9.0 g less, respectively, than Control-fed mice following 3 weeks of EAA restriction. Notably, Ile-D mice lost significantly more body weight than both Met-D and Leu-D mice, weighing on average 12.25 g less than controls at the study’s conclusion. The observed changes in body weight were largely independent of altered food intake patterns, with food intake being similar or higher in all groups eating an AA-depleted diet from the second week onwards (**Fig. 1C**). Together, these data suggest 3 weeks of individual EAA restriction is sufficient to induce whole body weight loss, likely via increased energy expenditure as has been previously reported in similar systems (^7,19,27,28^).

To assess changes in body weight as alterations in lean and fat mass stores, we analyzed body composition of each mouse at weeks 0 and 3. We found that only control mice retained lean mass over the course of our study. EAA-restricted mice experienced significant losses in lean mass that were greater than those induced by rapamycin treatment (**Fig. 1D**). Losses in lean mass correlated with a significant induction of autophagy in Leu-D mice as determined via decreased quadricep p62 protein expression (**Fig. S1A-B**). Although Leu-D has previously been shown to induce autophagy via mTORC1-inhibition, all mice were harvested following an overnight fast which may have masked autophagy induction relative to controls for Met-D and Ile-D treatment groups (^23^). In addition, EAA restriction induced significant losses in fat mass with a near total loss in detectable fat mass for Ile-D mice (**Fig. 1E**). Loss in fat mass for Leu-D and Ile-D mice was greater in magnitude than the observed losses in lean mass, resulting in an altered body composition consisting of increased lean mass and decreased fat mass percentages (**Figs. 1F-G**). The relative percentages of lean and fat mass were unchanged in Met-D mice, while control and rapamycin treated mice exhibited increased fat mass which is consistent with normal age-associated changes in body composition. Therefore, these data show dietary restriction of individual BCAAs has a greater impact on body composition than Met-D, likely via mTORC1-independent mechanisms.

In addition to altered body composition, dietary EAA restriction has been associated with improved glucose tolerance (^11–13)^. Although 3 weeks of complete dietary Met-D was unable to significantly improve glucose tolerance relative to controls, Leu-D and Ile-D mice exhibited improved glucose clearance under glucose tolerance test (GTT) conditions (**Figs. 1H-I**). Unlike EAA restrictions, rapamycin treated mice displayed reduced glucose tolerance as we have previously observed (^29,30^). Collectively, principal component analysis (PCA) of these data show 3 weeks of individual EAA (Met, Leu or Ile) dietary restriction is sufficient to induce whole-organism physiologic changes, with each EAA restriction eliciting distinct phenotypes which were not observed by rapamycin treatment alone (**Figs. 1J**).

### Specific dietary EAA restriction or rapamycin treatment induces distinct hepatic metabolite profiles

To determine how specific EAA restrictions influence cellular metabolism, we performed a targeted metabolomics analysis of livers from mice collected following the conclusion of the *in vivo* analysis above, after mice had consumed the indicated diets (or been treated with rapamycin) for 3 weeks (**Table S2**). Hierarchical clustering and principal component analysis (PCA) of treatment groups based on the altered relative abundance of over 100 quantifiable metabolites revealed changes induced by dietary EAA restriction were again more similar to one another than those stimulated by rapamycin treatment (**Figs. 2A-B**). However, unlike the grouping patterns observed in our physiologic measurements, Leu-D mice exhibited a metabolic profile which closely mimicked that of Met-D mice, while Ile-D fed animals were quite distinct from both Met-D and Leu-D-fed, as well as Control-fed animals. This result was largely unexpected as methionine, a non-BCAA, serves in dramatically different metabolic roles and biological pathways than do leucine or isoleucine.

**Figure 2:**
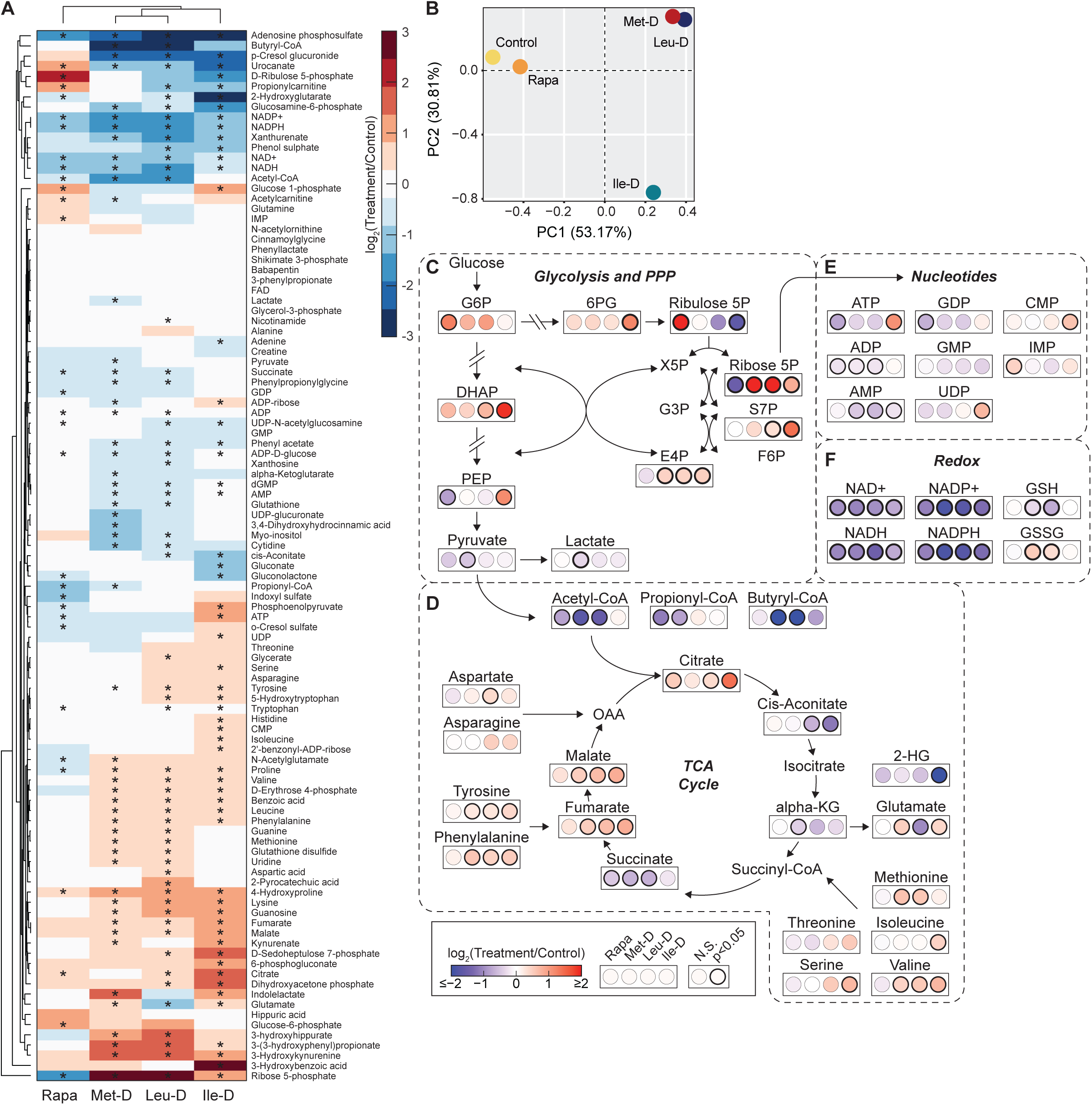
Alterations in hepatic metabolism following EAA restrictions and rapamycin treatment. (*A*) Heatmap with hierarchical clustering of metabolite abundance values. (*B*) PCA analysis of log transformed, relative metabolite abundance values. (*C-F*) Diagram depicting log2 fold-changes in metabolite abundance relative to controls within the larger context of their associated metabolic pathways. N = 12 for control, rapamycin, Met-D, and Ile-D groups; N = 11 for Leu-D; * = p-values measured using default MetaboAnalyst settings.

To assess how dietary restriction of EAAs altered specific metabolic pathways in the liver, we assessed the relative changes of key metabolites within and adjacent to glycolysis, the pentose phosphate pathway (PPP), the tricarboxylic acid (TCA) cycle, nucleotide, and redox metabolic networks (**Figs. 2C-F**). For glycolysis, a trending or significant increase in glucose 6-phosphate (G6P) and dihydroxyacetone phosphate (DHAP) was observed in all treatment groups although only Ile-D mice displayed an increased abundance of the downstream metabolite phosphoenolpyruvate (PEP) (**Fig. 2C**). Conversion of PEP to pyruvate appeared uniquely reduced in Met-D mice as pyruvate and lactate levels were both significantly lower than in control mice. Despite apparent EAA-specific effects on glycolytic metabolism, all treatment groups exhibited a trending or significant increase in all measured PPP intermediates with the exception of ribulose 5-phosphate (Ribulose 5P) which was largely unchanged in response to Met-D but significantly depleted in both Leu-D and Ile-D liver (**Fig. 2C**). Rapamycin elicited an opposing response in Ribulose 5P availability, with Ribulose 5P being significantly increased followed by a corresponding decrease in ribose 5-phosphate (Ribose 5P) (**Fig. 2C**). The metabolic changes induced by Rapamycin are consistent with known role for mTORC1 in positively regulating the translation of ribose-5-phosphate isomerase A, the enzyme that interconverts ribulose 5-phosphate to ribose 5-phosphate (^31^).

Downstream of glycolysis, all treatments besides Ile-D stimulated a decrease in acetyl-CoA levels relative to controls (**Fig. 2D**). Similar trends were observed for propionyl- and butyryl-CoA, highlighting a general decrease in liver short chain acyl-CoA levels for Met-D, Leu-D, and rapamycin treated mice. Despite decreased acetyl-CoA availability for Met-D, Leu-D, and rapamycin treated mice, all treatment groups exhibited trending or significant increases in citrate availability (**Fig. 2D**). However, elevated citrate levels were not associated with a greater abundance of immediate downstream TCA cycle intermediates. Instead, all conditions exhibited similar or decreased abundance of cis-aconitate, alpha-ketoglutarate (alpha-KG), and/or succinate relative to controls. 2-hydroxyglutarate (2-HG), a metabolite with structural similarities to alpha-KG, was also significantly decreased in Ile-D mice (**Fig. 2D**). A significant increase in fumarate and malate under Met-D, Leu-D, and Ile-D, but not rapamycin treatment, mirrored increases in tyrosine and phenylalanine abundance, suggesting the increase in malate/fumarate is driven by selective amino acid anaplerosis (**Fig. 2D**). Along with tyrosine and phenylalanine, numerous other amino acids were generally elevated in response to each dietary EAA restriction, but not rapamycin treatment (**Figs. 2A and 2D**). In fact, the EAA designated for dietary restriction uniformly presented with elevated hepatic levels. We have previously observed that 5 weeks of Met-D is sufficient to lower liver methionine levels >50% (^22^), while the current study of dietary EAA restriction (for 3 weeks) shows an increase in liver methionine, likely reflecting the window of adaptive response where methionine is salvaged from other tissues.

In addition to central metabolic pathways, other major metabolites showed differential regulation in response to dietary EAA restrictions. For example, Ile-D led to the accumulation of high energy state adenosine and guanosine nucleotides (i.e, ATP and GDP) which showed a trending or significant decrease across the other treatment groups (**Fig. 2E**). Uridine, cytidine, and inosine nucleotides were also elevated in Ile-D mice with no clear pattern observed in response to other dietary EAA restrictions or rapamycin treatment. Interestingly, the redox metabolites NAD^+^/NADH and NADP^+^/NADPH were significantly reduced across all conditions, including rapamycin treatment, suggestive of an overall decrease in NAD^+^ synthesis and/or increase in NAD^+^ catabolism (**Fig. 2F**). A decreased ratio of glutathione (GSH) to glutathione disulfide (GSSG) was observed in Met-D and Leu-D mice with no changes in the relative abundance of either redox metabolite altered in Ile-D or rapamycin mice. Together, these metabolic analyses show dietary Ile-D has a unique impact on the relative levels of major metabolic cofactors with dietary Met-D and Leu-D displaying common responses unique from both Ile-D and rapamycin-treated mice.

### Dietary EAA restrictions stimulate unique Histone PTM patterns across all conditions that are not mimicked by mTORC1 inhibition

Fluctuations in metabolite availability can regulate cellular processes through numerous, diverse mechanisms. This includes direct regulation of the epigenome, as the enzymes responsible for adding and removing epigenetic post-translational modifications (PTMs) rely on metabolic co-substrates to support their catalytic activity. Notably, all three EAAs investigated in this study are metabolized to such co-substrates. To determine whether the liver epigenome is uniquely sensitive to dietary EAA restrictions or rapamycin treatment, we performed LC-MS/MS histone proteomics on liver tissue harvested at the study’s completion (**Table S3**). Hierarchical clustering and PCA analyses based on the altered relative abundance of 93 uniquely modified histone peptides revealed distinct global histone PTM profiles for all treatment groups (**Figs. 3A-B**). We previously investigated the epigenetic responses to Met-D in detail, revealing adaptive chromatin mechanisms highlighted by preferential H3K9me1 maintenance in the presence of global decreases to higher-state histone methylations (^22,32^). These trends under Met-D were replicated in the present study (**Fig. 3C-E**).

**Figure 3:**
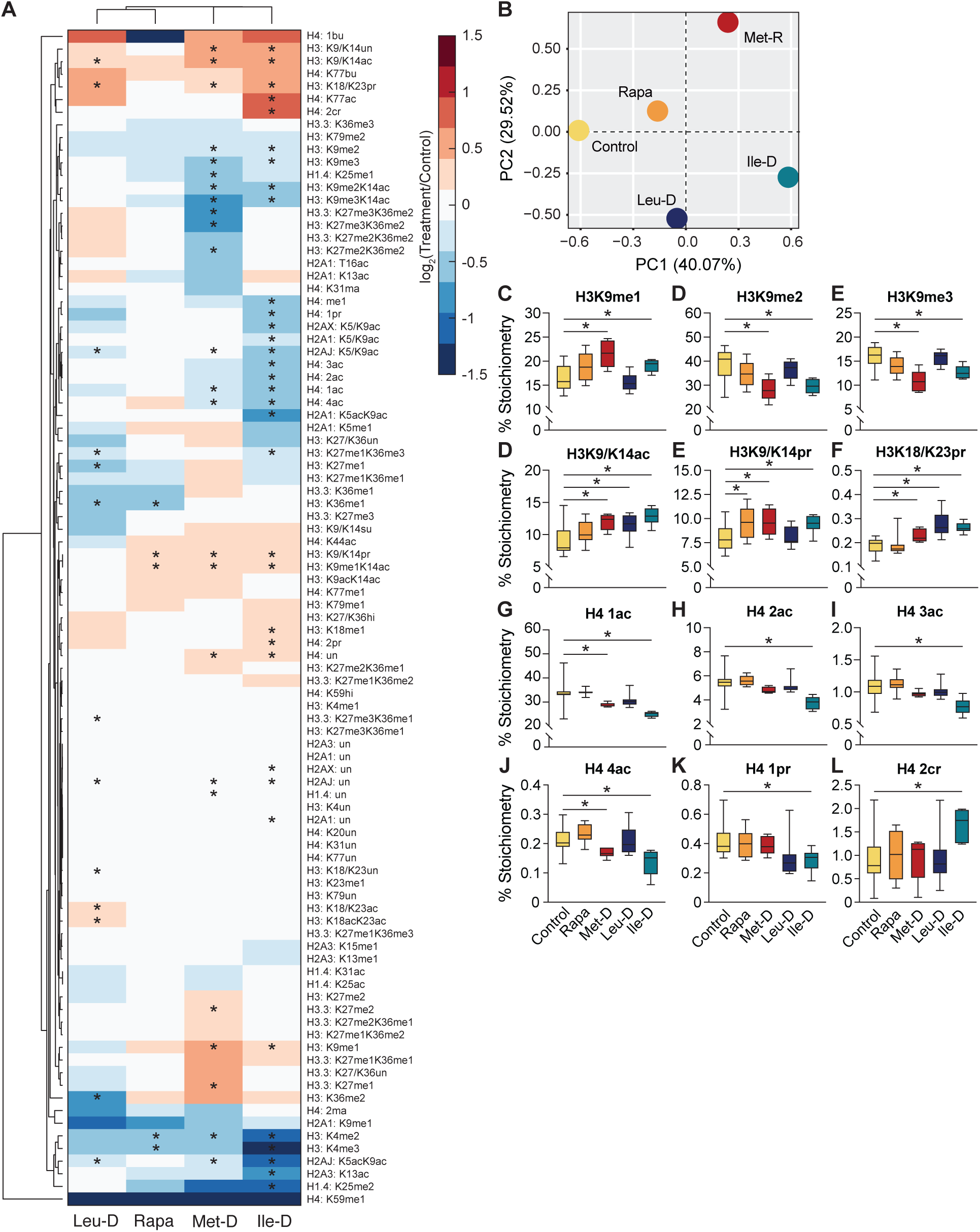
Hepatic histone PTM responses to EAA restrictions and rapamycin treatment. (*A*) Heatmap with hierarchical clustering of histone peptide stoichiometric abundance values compared to control values. (*B*) PCA analysis of log transformed, relative histone peptide stoichiometry values. (*C-L*) Box plots depicting deconvoluted stoichiometry percentages of residue-specific histone PTMs. Error bars = standard deviation; N = 12 for control, N = 10 for Rapa; N = 7 for Leu-D; N = 6 for Met-D and Ile-D; * = p-value < 0.05 as measured via Student’s t-test.

Unexpectedly, Ile-D and Met-D groups exhibited a highly similar trend in H3K9 methylation states, as well as increased acetylation on H3K9/K14 and increased propionylation on both H3K9/K14 and H3K18/K23 (**Figs. 3C-F**). All three EAA restrictions led to similar increases in H3 acylation states, but rapamycin treatment only stimulated a significant increase in H3K9/K14 propionylation. Interestingly, unlike the increase of H3 acylations, Met-D and Ile-D led to significantly decreased levels of acetylation on H4 (**Fig. 3G-J**). The decrease in H4 acetylation was especially prevalent in Ile-D liver as all acetylated H4 peptide species (i.e., 1ac, 2ac, 3ac, and 4ac) were reduced under this condition. While Met-D mice displayed decreased liver acetyl-CoA levels, Ile- D acetyl-CoA levels were unchanged. These data either suggest independent mechanisms dictate H4ac profiles in response to dietary Met-D/Ile-D, or that H4 acetylation mechanisms are shared but function independent of whole-cell acetyl-CoA availability. In support of the latter, H4 acetylation levels in Leu-D and rapamycin treated mice were robust despite decreased liver acetyl-CoA availability (**Fig. 3G-J**). Notably, unlike acetylation levels on H4, other short chain acylations were largely unchanged across conditions with only Ile-D mice exhibiting slight increases in butyrylated and crotonylated H4 residues (**Fig. 3K-L**). Together, these data demonstrate that 3-week dietary EAA restrictions or rapamycin treatment have differential effects on the liver epigenome, providing a potential mechanism driving EAA restriction-specific phenotypes.

### Ile-D induces a robust transcriptional response compared to Met-D, Leu-D, or Rapa treatment

To investigate how each treatment altered the functional gene expression landscape of the liver, we performed bulk RNA-sequencing (**Table S4**). PCA of log transformed gene counts per million (CPM) values revealed identical groupings to those observed from the metabolomics analyses (**Figure 4A**), suggesting a direct functional link between the changes in transcripts and subsequent metabolic pathways. Also, the similar PCA groupings of Met-D and Leu-D further suggest that these amino acid deficiencies are sensed through a common mechanism, distinct from Ile-D. In support of this observation, 45.6% (n = 4088) of all differentially expressed (DE) genes identified in Ile-D liver were uniquely altered under this condition alone (**Fig. 4B**). Only 11.7% (n=590) and 13.2% (n=494) of DE genes in Leu-D and Met-D mice, respectively, were unique to each given condition (**Fig. 4B**). In fact, a majority of Leu-D or Met-D DE genes were also differentially regulated by at least one other dietary EAA restriction (i.e., Leu-D = 83%, Met-D = 81.4%). A majority of rapamycin DE genes (77.5%) were also shared by at least one other EAA restricted condition, although relative to the changes observed in other conditions, the absolute number of DE genes contributing to this pattern (n=478) was significantly lower (**Fig. 4B**). The absolute numbers of identified DE genes mirrored the relative magnitude of each response, with Met-D and Leu-D DE genes eliciting transcriptional responses less significant than Ile-D but greater than rapamycin treatment (**Figs. 4C-4F**). Together, these data collectively suggest EAA restrictions stimulate distinct hepatic transcriptional responses from rapamycin treatment.

**Figure 4:**
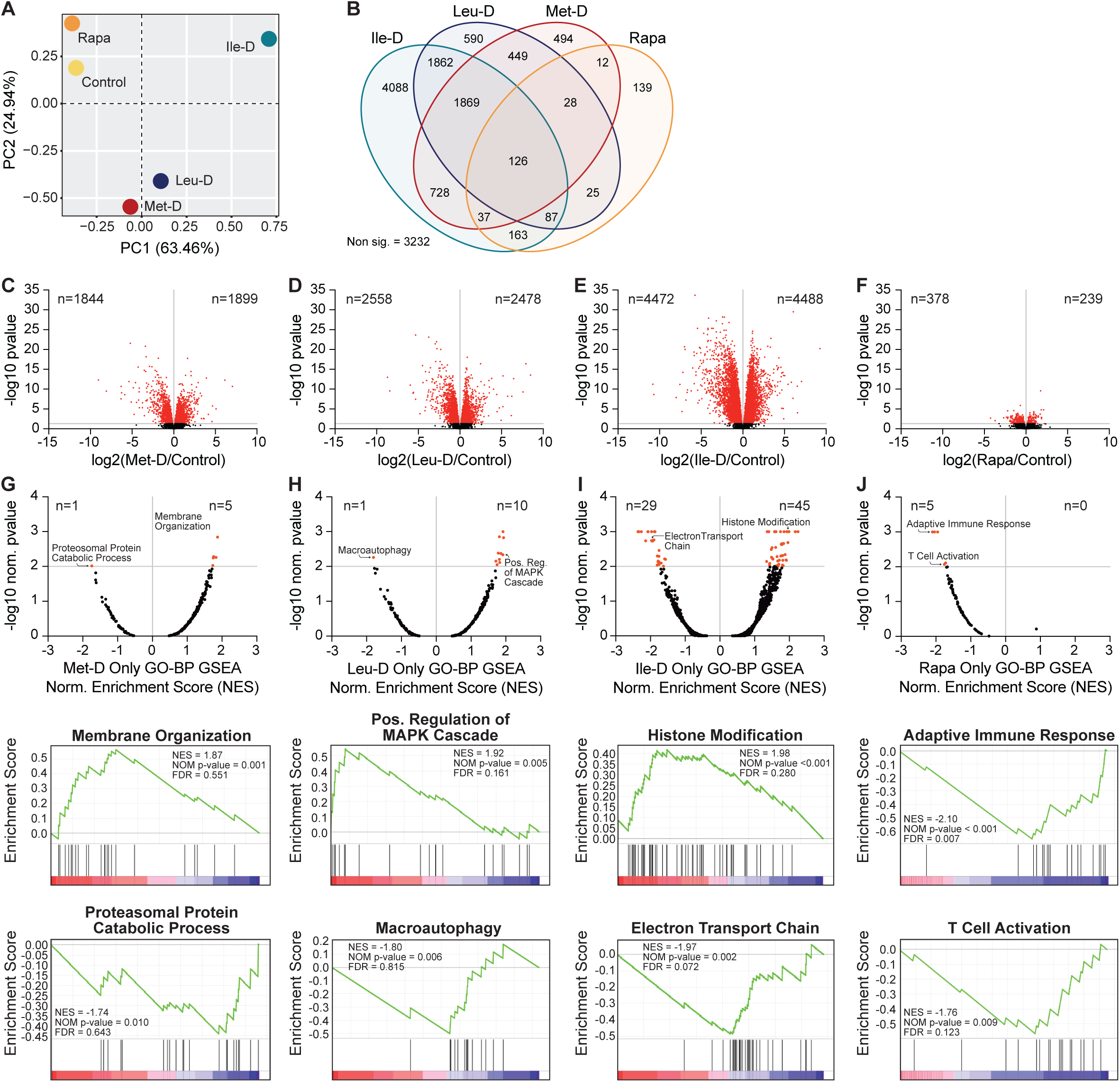
Hepatic transcriptional programs elicited by EAA restrictions and rapamycin treatment. (*A*) PCA analysis of log transformed gene CPM values. (*B*) Venn diagram of differentially expressed genes with p<0.05. (*C-F*) Volcano plots depicting log2 FC and -log10 adj. p-values for all identified genes. Horizontal marker indicates threshold for statistical significance (p-adj. < 0.05) with significantly changing genes marked in red. (G-J) Dot plots illustrating enriched Gene Ontology – Biological Process (GO-BP) gene sets as determined via Gene Set Enrichment Analyses (GSEA) of differentially expressed genes unique to each given condition. Example GSEA plots highlighting significantly enriched GO-BP gene groups with positive and negative normalized enrichment scores (NES) are present beneath each corresponding dot plot. N = 12.

With EAA transcriptional responses being clearly distinct from those elicited by rapamycin treatment, we performed Gene Ontology Biological Process (GO-BP) functional annotation of DE genes shared by Met-D, Leu-D, and Ile-D conditions to identify candidate processes which are broadly responsive to dietary EAA restriction (^33^). This analysis identified 59 significantly enriched GO-BP terms covering cellular functions related to lipid (e.g., fatty acid and cholesterol metabolic processes) and amino acid (e.g., glutathione and glutamate metabolic processes) metabolism as well as chromatin regulation (e.g., chromatin remodeling and chromatin looping) and RNA processing (e.g., rRNA processing and RNA splicing) (**Table S5**). Notably, the GO-BP term “transmembrane transport” possessed the second-most significant FDR value and was primarily enriched for solute carriers (SLCs), including the amino acid transporters SLC25A22 and SLC36A4. These results highlight how dietary EAA restrictions uniformly stimulate significant responses by nutrient uptake and utilization pathways in addition to shared aspects of chromatin and RNA regulation.

We next sought to assess the unique functional consequences of each individual EAA restriction or rapamycin treatment by analyzing condition-specific DE genes via a GO-BP pre-ranked Gene Set Enrichment Analysis (GSEA) (**Figs. 4G-J, Table S6**) (^34^). Met-D-specific DE genes were assigned to a small number of significantly enriched gene sets (n=6) which primarily possessed positive NES values and consisted of membrane organization/cell motility biological processes (**Fig. 4G**). A similar number of significantly enriched gene sets were identified using Ile-D specific transcripts (n=11) which also predominantly held positive NES scores. Positive NES gene sets were overwhelmingly associated with diverse intracellular signal transduction pathways such as the MAPK signaling cascade (**Fig. 4I**). Given the greater magnitude of transcriptional response identified in Ile-D mice, it was unsurprising to find that Ile-D-specific DE genes were significantly enriched in the greatest number of GO-BP gene sets (n=74) (**Fig. 4J**). Interestingly, multiple positive NES gene sets were associated with histone/protein post-translational modifications, providing potential insights into mechanisms regulating the unique histone PTM profile of Ile-D mice (**Fig. 3A**). Negative NES gene sets were primarily associated with lipid metabolism and mitochondrial oxidative phosphorylation, suggestive of a compensatory mechanism aimed towards preserving liver lipid stores in the presence of a near complete loss in total fat mass(**Fig. 1E**). Lastly, as observed in both metabolomics and histone proteomics datasets, rapamycin-specific enriched gene sets (n=5) were again distinct from all dietary EAA restrictions and primarily consisted of NES gene sets related to immunity-associated transcriptional programs (**Fig. 4G**). Therefore, these data collectively reveal that Ile-D stimulates the largest and most distinct transcriptional response of all treatments while providing insight into underlying mechanisms supporting this intervention’s unique physiologic, metabolic, and epigenetic phenotypes.

### Multi-Omics Factor Analysis identifies differential histone acetylation and expression of central metabolic genes as defining features of Ile-D

To generate a quantitative understanding of the molecular responses unique to Ile-D, a Multi-Omics Factor Analysis (MOFA) was performed (**Extended Data Figure 2**). MOFA is an unsupervised computational method that co-analyzes datasets to identify a set of Factors which explain the biological and technical variability between experimental groups (^35^). When applying MOFA to the 14,139 data points collected in this study, 15 distinct factors were identified with a majority of the variability explained by Factors 1 through 3 (**Fig. 5A**). Ile-D measurements strongly correlated with Factor 1 (r= 0.9) whereas remaining treatment groups possessed either negligible (Leu-D: r= 0.06, Met-D: r= -0.14) or negative correlations with this Factor (Control: r= -0.4, Rapa: r= - 0.41) (**Fig. 5B**). The percent of total variance contributed by each dataset was greatest in Factor 1, supporting our observation that Ile-D elicits distinct responses across all measurements taken in this study (**Fig. 5A**). Additionally, Met-D and Leu-D treatment groups strongly correlated with Factor 2, further suggesting dietary methionine or leucine restriction may function through similar mechanisms.

**Figure 5:**
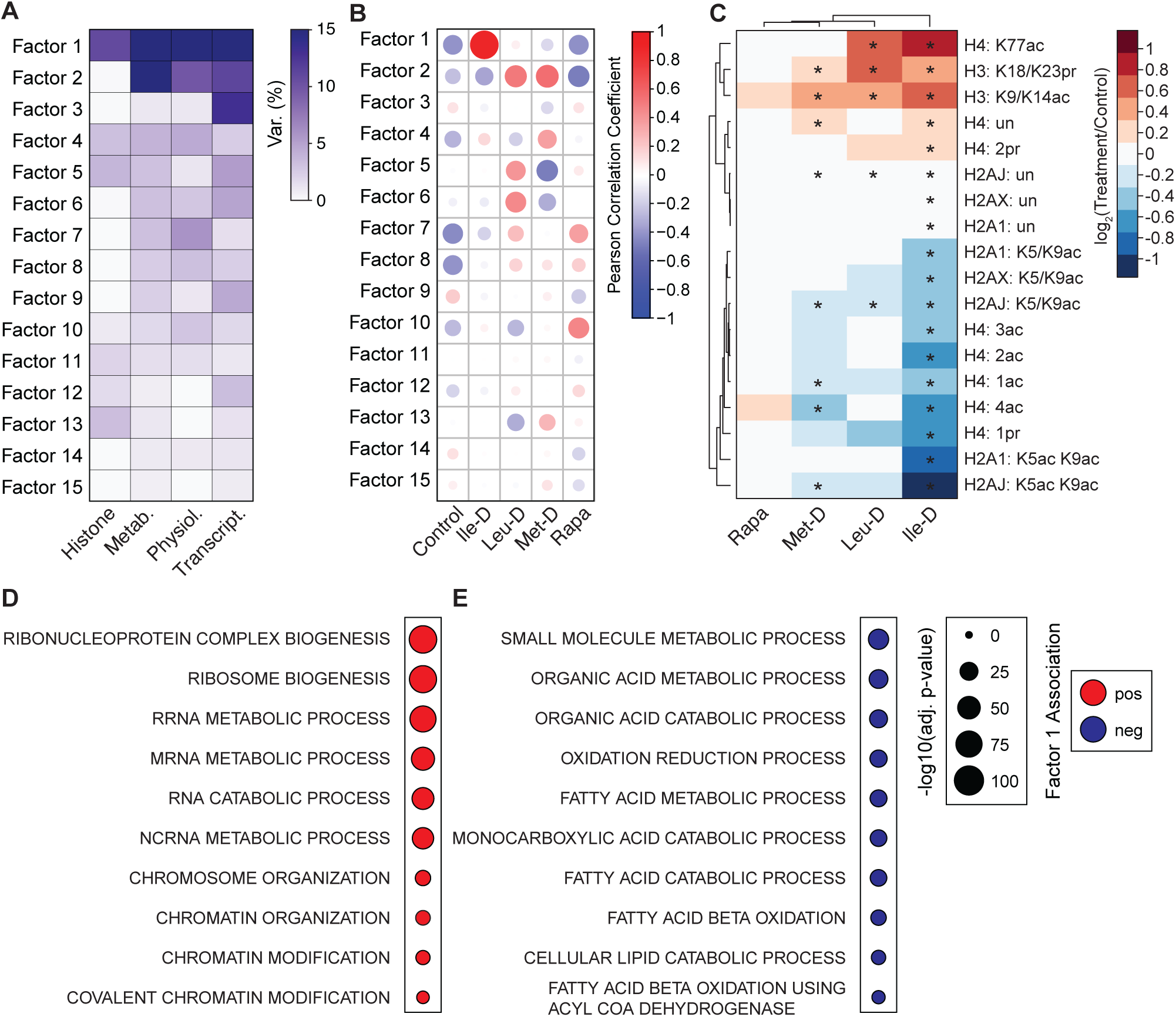
Multi-Omics Factor Analysis of whole organism and hepatic molecular measurements. (*A*) Heatmap of variance decomposition by factor. (*B*) Plot illustrating Pearson correlation coefficients for control and treatment groups with 15 MOFA Factors. (*C*) Heatmap with hierarchical clustering of Factor 1-associated histone peptide stoichiometric abundance values compared to control values. Factor 1-associated histone peptides were determined using a weight cutoff value of +/- 0.2 which identified the top 18 associated histone peptides. (*D-E*) Plots depicting a subset of significantly enriched gene sets identified using Factor 1 transcripts. All significantly enriched gene sets can be found in Extended Data Figure 2. * = p-value < 0.05 as measured via Student’s t-test.

Interestingly, the variance contributed by histone PTM measurements was primarily associated with Factor 1 (10.84%), suggesting differences in histone PTM regulation most strongly distinguish Ile-D from remaining treatment groups (**Fig. 5A**). When assessing which histone PTMs assigned to Factor 1 primarily contributed to this variance, we detected a redistribution in histone acetylation across H3, H4, and H2A isoforms (**Fig. 5C**). Specifically, we identified hypoacetylation at H4 K5/K8/K12/K16 and H2A1/J/X K5/K9 with hyperacetylation at H3 K9/K14 and H4 K77ac (**Fig. 5C**). This pattern of acetyl-group redistribution is supported by the GSEA results generated from Ile-D-specific DE genes which identified the H3K14 acetyltransferase Kat7 as well as the broad histone deacetylases Sirt1 and Hdac4 in the “Histone Modification” gene set (**Fig. 4J**). Together, these MOFA Factor 1-associated data suggest differential regulation of histone acetylation profiles is a unique and defining characteristic of Ile-D in the liver.

As the epigenome directly regulates gene expression profiles, we next analyzed Factor 1 Principal Component Gene Set Enrichment (PCGSE) results to identify which pathways are transcriptionally associated with this redistribution in histone acetylation. PCGSE of Factor 1 genes identified an upregulation of pathways linked with RNA metabolism and processing as well as chromatin organization and modification (**Fig. 5D**). Notably, “Chromatin Modification,” “Chromatin Organization,” and “Covalent Chromatin Modification” gene sets included Kat7, Sirt1, and Hdac4, further implicating these chromatin modifiers as mediators of the epigenetic response to Ile-D. Downregulated Factor 1 gene sets were primarily enriched for organic acid metabolism and lipid oxidation, two pathways critical for mitochondrial acetyl-CoA production (**Fig. 5E**). The robust transcriptional downregulation of mitochondrial acetyl-CoA producing pathways were striking as hepatic acetyl-CoA levels were maintained during Ile-D, suggesting alternative pathways may be engaged to support acetyl-CoA maintenance under this condition. Together, these MOFA PCGSE results highlight the association of chromatin regulation and mitochondrial acetyl-CoA metabolism with histone acetylation redistribution in Ile-D liver.

### Ile-D induced metabolic and epigenetic remodeling occur rapidly within isolated cells

Although distinct acetyl-CoA metabolism and histone acetylation states are unique features of Ile-D within the liver, it remains unclear whether these mechanisms require complex inter-cellular/organ signaling or represent inherent, fundamental intracellular responses. To determine whether isolated cells possess the inherent capability of initiating these molecular responses, we investigated the metabolic and epigenetic responses of human liver HepG2 cells over 24 hours of Ile-D (**Fig. 6A, Tables S7 and S8**). *In vitro*, rapid removal of Ile from culture media led to a dramatic loss in its downstream catabolic product KMVA, which was no longer detectable after 10 minutes (**Figs. 6B-C**). Importantly, this perturbation had minimal effects on global BCAA metabolism as leucine, valine, and their metabolic products KICA and KIVA, respectively, exhibited minor increases in abundance (**Figs. 6B-C**). Similarly to the *in vivo* metabolomics data, Ile-D HepG2 cells were largely capable of maintaining cellular acetyl-CoA pools (**Fig. 6D**). This maintenance of acetyl-CoA was particularly striking as propionyl-CoA and succinyl-CoA, downstream acyl-CoA products of isoleucine metabolism, were rapidly depleted and maintained at low levels over 24 hours of Ile-D (**Fig. 6D**).

**Figure 6:**
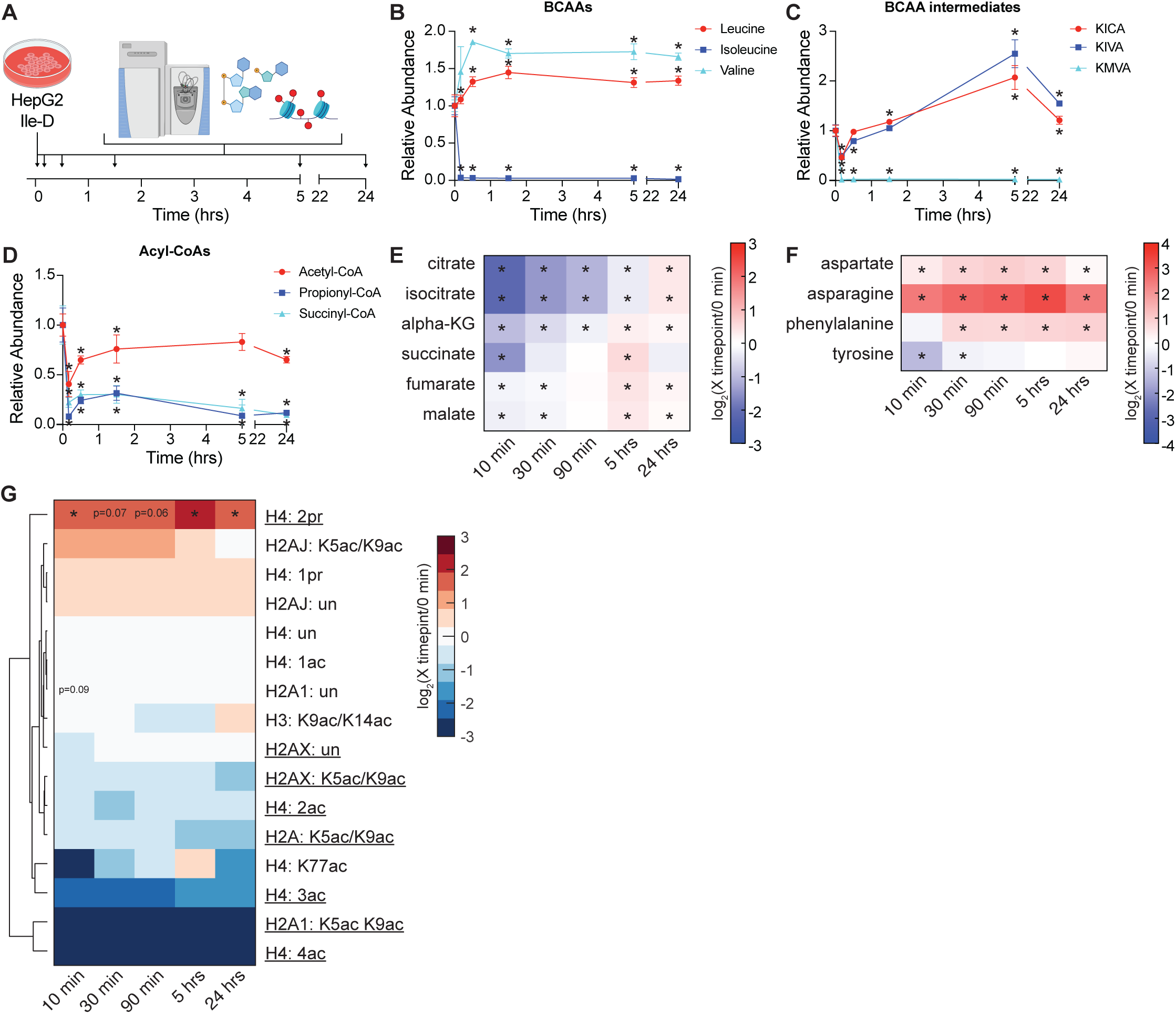
Metabolic and epigenetic mechanisms elicited by Ile-D of isolated cells. (*A*) Schematic depicting experiment design for *in vitro* Ile-D of HepG2 hepatocellular carcinoma cells. (*B-C*) Plots depicting changes in metabolite abundance relative to 0 hr control values. (*E-F*) Heatmaps of metabolite abundance values. (*G*) Heatmap with hierarchical clustering of histone peptide stoichiometric abundance values compared to 0 hr control values. Depicted peptides of interest were originally identified via MOFA in Fig 5. Histone peptides with similar changes in abundance *in vitro* as previously identified *in vivo* are underlined. Error bars = STDEV; N = 4 for metabolite measurements; N = 3 for histone PTM measurements; * = p-value < 0.05 as measured via Student’s t-test.

*In vivo*, maintained acetyl-CoA levels were accompanied by selective amino acid anaplerosis which restored TCA cycle intermediate abundance from fumarate to citrate in contrast to the decreased abundance of intermediates from cis-aconitate to succinate (**Fig. 2D**). *In vitro,* Ile-D induced similar changes in TCA cycle intermediate abundance, with alpha-KG and succinate both significantly decreased over the initial 30 minutes (**Fig. 6E**). Fumarate and malate were generally maintained during this period although the abundance of both metabolites decreased slightly (**Fig. 6E**). Notably, the amino acids aspartate, asparagine, and phenylalanine all increased in abundance over the initial 30 minutes of Ile-D, which again implicates selective amino acid anaplerosis in supporting fumarate and malate levels (**Fig. 6F**). However, the relative abundances of all TCA cycle intermediates recovered to or above pre-restriction levels by 24 hours, coinciding with maintained elevation of anaplerotic amino acids, revealing that isolated HepG2 cells are capable of restoring TCA cycle metabolism under prolonged Ile-D *in vitro* (**Fig. 6E-F**). These data indicate TCA cycle metabolism and selective anaplerosis are similarly responsive to Ile-D both *in vitro* and *in vivo*, with the immediate *in vitro* response (*i.e.*, 10-30 min) being most similar to 3-week Ile-D mouse liver.

In addition to possessing similar metabolic responses to *in vivo* Ile-D, HepG2 cells also exhibited a similar reorganization of histone acetylation (**Fig 5A**). Specifically, Ile-D HepG2 cells displayed hypoacetylation at H4 K5/K8/K12/K16 and H2A/X K5/K9 and maintained levels of H3 K9/K14ac (**Fig. 6G**). Changes in histone PTM abundance were established during the first 10 minutes of Ile-D and held largely constant over the entire 24-hour time course (**Fig. 6G**). Together, these data illustrate that Ile-D stimulated remodeling of acetyl-CoA and TCA cycle metabolism, as well as unique regulation of histone acetylation, are inherent, fundamental responses of isolated cells.

## Discussion

In this study, we set out to gain new insight into the molecular mechanisms by which dietary EAA availability regulates physiologic and hepatic responses at the molecular level. We found that individual depletion of EAAs led to both common and diet-specific changes. Consistent with prior work, individual depletion of Ile, Leu, or Met led to improved glucose tolerance with losses in fat and lean mass (^12,22,36,37^). Here, EAA restriction-induced changes in glucose tolerance and body composition were accompanied by dynamic alterations to the hepatic metabolome, epigenome, and transcriptome. Metabolically, increases in hepatic glycolytic and PPP intermediates were prominent across EAA restrictions. It has been estimated that as much as ⅓ of incoming glucose can be consumed by the PPP (^38,39^), suggesting increased glucose flux through the PPP pathway could promote greater glucose clearance. An increased abundance of free amino acids in the liver, even among the amino acids being restricted, was identified across all restrictions. An increase in hepatic amino acid levels, which was accompanied by a progressive decline in lean mass, is consistent with skeletal muscle loss (autophagy) and the freeing up of AAs for essential protein synthesis and other metabolic processes throughout the body. Together, these data highlight the trend of liver self-preservation in response to EAA restrictions at the expense of skeletal muscle and fat stores.

While a subset of physiologic and hepatic molecular responses was shared across EAA restrictions, unique response patterns were identified. For example, we were surprised to find that leucine and methionine restriction stimulated highly similar effects in liver metabolite and gene expression programs. Multi-omics factor analysis (MOFA) objectively identified the similarity between Leu-D and Met-D molecular responses, uniquely correlating both treatment groups with Factor 2, a Factor with high variance contributions from hepatic metabolite and transcript measurements (**Fig. 5A-B**). Factor 2 metabolite trends included a general accumulation of amino acids, which corresponded with the apparent downregulation of amino acid catabolism genes. Decreased acetyl-CoA abundance was also associated with Factor 2, highlighting the disparate mechanisms by which Met-D/Leu- D and Ile-D regulate acetyl-CoA metabolism.

Unique from Met-D and Leu-D, Ile-D consistently elicited distinct responses across all physiological and molecular measurements taken. This distinct response was confirmed via MOFA, which uniquely correlated Ile- D with Factor 1-associated features. PCGSE of Factor 1 genes identified an upregulation of chromatin regulation as well as the downregulation of organic acid metabolism and lipid oxidation pathways. This latter difference between Ile-D animals and Met-D/Leu-D animals was unexpected as the Ile-D liver was uniquely able to maintain acetyl-CoA levels. These data suggest acetyl-CoA is likely maintained through alternative pathways under Ile-D. Our metabolomics analysis implicates cytoplasmic acetyl-CoA synthesis via ACLY as one candidate pathway. Citrate, ACLY’s obligatory substrate, shows the greatest magnitude of accumulation in Ile-D liver. Additionally, our metabolomics data suggest there is a significant increase in TCA cycle anaplerosis via the conversion of phenylalanine and tyrosine to fumarate, which would provide additional carbons for acetyl-CoA and citrate synthesis. The enzyme responsible for catalyzing the first step of tyrosine catabolism, TAT, is upregulated in Ile- D liver (**Table S5**). Importantly, tyrosine and phenylalanine breakdown through the TAT pathway generates acetoacetate, which can be further metabolized to two molecules of acetyl-CoA independent of ACLY activity (^40^). Very recent work has shown that acetoacetate-derived acetyl-CoA is capable of supporting *de novo* lipogenesis upon inhibition of the mitochondrial citrate shuttle, highlighting the flexibility of cells to generate cellular acetyl-CoA through alternative pathways when necessary (^41^). Further work will be needed to demonstrate that carbon flux through Phe/Tyr fuels acetyl-CoA synthesis when adipose tissue is depleted, as in Ile-D.

Although acetyl-CoA levels were maintained, the significant redistribution of histone acetylation was identified as a discriminating feature of Ile-D liver. This included the global hypoacetylation of H2A and H4 with a corresponding increase in H3K9/K14 acetylation. The opposing trends in H4 and H3 acetylation are consistent with a previous report in which Hat1 acetylated, nascently translated H4 proteins are subsequently deacetylated following their transport to the nucleus. The liberated acetyl-groups were then utilized to provide a local co- substrate supply and support targeted H3K9 acetylation (^42^). Here, consistent with this mechanism, Hat1 and the histone deacetylases Hdac4 and SIRT1 are specifically upregulated in response to Ile-D. It is interesting to speculate that more efficient utilization of acetyl-CoA in response to Ile-D functions in cooperation with selective anaplerosis, acetoacetate catabolism, and/or ACLY-dependent acetyl-CoA production to maintain global metabolite pools under this condition. If true, such a cooperative mechanism would be required to maintain global acetyl-CoA availability in the presence downregulated amino acid catabolism, fatty acid metabolism, and organic acid metabolism as acetyl groups stored on histone proteins are not present in adequate amounts to support total cellular requirements (^43^). However, H4 deacetylation has been shown to liberate acetyl groups for utilization outside the nucleus when needed (^44^). Notably, Ile-D HepG2 cells were also largely capable of maintaining acetyl- CoA levels. This corresponded with a remodeling of amino acid and TCA cycle metabolism as well as differential acetyl-CoA utilization by histone acetyltransferases, closely mimicking our *in vivo* observations. Therefore, these data suggest the ability to cooperatively remodel acetyl-CoA metabolism and histone acetylation profiles is an inherent capability of isolated cells under Ile-D.

In this study, we also assessed the ability of mTORC1 inhibition via rapamycin to replicate the physiologic and molecular responses to EAA restrictions. Overall, we found minimal overlap among our hepatic metabolomic, epigenetic, and transcriptomic analyses. In fact, rapamycin treatment elicited relatively muted responses compared to dietary EAA restrictions across all physiologic and molecular responses assayed in this study. Together, these observations imply that downregulation of the mTORC1 signaling pathway in response to select EAA restrictions plays a minor role in facilitating whole organism and hepatic molecular responses. However, a potential limitation of this conclusion is that there are rapamycin-resistant functions of mTORC1 that could be engaged by EAA restriction (^45^), making additional work with genetically engineered mouse models of altered mTORC1 signaling needed to confirm that EAAs largely function through mTORC1-independent pathways. Our study is also limited by its focused analysis of male C57BL/6J mice, as rapamycin and dietary EAA restrictions are known to elicit sex-specific phenotypes (^46,47^). Similarly, although all molecular measurements in this study focused on hepatic responses, the inherent ability of isolated cells to mount a conserved response to Ile-D was limited to the analysis of a singular, non-primary liver cell line. Future studies are also needed to assess how the molecular responses to individual EAA restrictions identified here can be stimulated by low, but still physiologic levels of EAA restriction and to determine the time-dependence of which these diets influence physiologic and molecular phenotypes.

In conclusion, we have shown that dietary restriction of single EAAs, while having similar effects on glucose homeostasis, body composition, and general amino acid catabolism, can yield individual responses by the hepatic metabolome, epigenome, and transcriptome which are independent of mTORC1 activity. Notably, Ile-D elicited the most significant and unique response, defined by distinct anaplerosis, decreased BCAA and fatty acid catabolism, increased pentose phosphate pathway and Phe/Tyr catabolism, as well as histone H4/H2A hypoacetylation and histone H3K9/K14 hyperacetylation. Together, these results highlight the potential importance of dietary protein EAA composition in regulating metabolic health at the molecular level.

## Methods

### Animals and Diets

All procedures were performed in conformance with institutional guidelines and were approved by the Institutional Animal Care and Use Committee of the William S. Middleton Memorial Veterans Hospital (Madison, WI, USA). Male C57BL/6J (#000664) mice were obtained from The Jackson Laboratory. All mice were housed three per cage, maintained at a temperature of approximately 22°C, and health checks were completed on all mice daily. Studies were commenced when the mice were 14 weeks of age.

Mice were then randomized to receive the indicated diets from Inotiv either the Control (TD.01084), Met-D (TD.140119), Leu-D (TD.180789) or Ile-D (TD.190879) diets, or the Control diet plus rapamycin (delivered I.P. in 5% Tween 20, 5% PEG40, 0.9% NaCl, 3% ethanol at 4 mg/kg/day). Full diet descriptions are provided in **Table S1**. The randomization of mice was performed at the cage level to ensure that all groups had approximately the same initial starting weight and body composition. Mice were housed in a SPF mouse facility in static microisolator cages, except when temporarily housed in a Columbus Instruments Oxymax/CLAMS-HC metabolic chamber system. Mice were housed under a 12:12 h light/dark cycle with free access to food and water, except where noted in the procedures below.

### In vitro Cell Culture

Human HepG2 cell lines were initially cultured in EMEM media (Corning, 10-009-CV) containing all amino acids and 10% FBS at 37°C, 5% CO_2_, while EMEM media devoid of amino acids (US Biological, M3859-01) was supplemented with all amino acids except isoleucine to generate isoleucine depleted media for the appropriate studies. All non-isoleucine amino acids were added according to their replete EMEM concentrations. Cells were seeded at a seeding density of 0.4 x10^6^ and 3 x10^6^ in 6-well plates and 100mm culture dishes for LC-MS metabolite analysis and histone proteomics respectively. Prior to Ile depletion, complete EMEM media was replaced with Ile-Replete media, supplemented with 10% dialyzed FBS (Gibco, A3382001) for one hour. This was followed by rinsing the cells with PBS, pH 7.4, and subsequent addition of Ile-Deplete media (with 10% dialyzed FBS) for restriction durations of 10 min, 30 min, 1.5 hr, 5 hr, and 24 hr. At the end of each restriction time point, cells were harvested for metabolites and core histones extraction accordingly.

### In vivo Procedures

Glucose tolerance tests were performed by fasting the mice overnight for 16 hours and then injecting glucose (1 g kg^−1^) (^7,48^). Glucose measurements were taken using a Bayer Contour blood glucose meter (Bayer, Leverkusen, Germany) and test strips. Mouse body composition was determined using an EchoMRI Body Composition Analyzer (EchoMRI, Houston, TX, USA). For assay of multiple metabolic parameters [O_2_, CO_2_, food consumption, respiratory exchange ratio (RER), energy expenditure] and activity tracking, mice were acclimated to housing in a Oxymax/CLAMS-HC metabolic chamber system (Columbus Instruments) for ∼24 h and data from a continuous 24 h period was then recorded and analyzed. Food consumption in home cages was measured by moving mice to clean cages, filling the hopper with a measured quantity of fresh diet in the morning and measuring the remainder in the morning 3 days later. The amount was adjusted for the number of mice per cage, the number of days that passed and the relative weights of the mice (i.e., heavier mice were credited with a larger relative portion of the food intake). Mice were euthanized by cervical dislocation after an overnight (16h) fast and tissues for molecular analysis were flash-frozen in liquid nitrogen and stored at -80°C.

### Immunoblotting

Tissue samples from muscle (quadriceps, lateral, on femur) were lysed in cold RIPA buffer supplemented with phosphatase inhibitor and protease inhibitor cocktail tablets (Thermo Fisher Scientific, Waltham, MA, USA) using a FastPrep 24 (M.P. Biomedicals, Santa Ana, CA, USA) with bead-beating tubes (16466–042) from (VWR, Radnor, PA, USA) and zirconium ceramic oxide bulk beads (15340159) from (Thermo Fisher Scientific, Waltham, MA, USA). Protein lysates were then centrifuged at 13,300 rpm for 10 min and the supernatant was collected. Protein concentration was determined by Bradford (Pierce Biotechnology, Waltham, MA, USA). 24 μg protein was separated by SDS– PAGE (sodium dodecyl sulfate–polyacrylamide gel electrophoresis) on 10% resolving gels (ThermoFisher Scientific, Waltham, MA, USA) and transferred to PVDF membrane (EMD Millipore, Burlington, MA, USA). Imaging was performed using the BioRad Chemidoc Imaging System (primary antibodies: SQSTM1/p62 (1:1000; American Research Products, #03-GP62-C; host: Rabbit) and GAPDH (1:1000; Cell Signaling Technology, #5174; host: Guinea Pig). secondary antibodies: Anti-Guinea pig (1:2000; American Research Products, #90001; host: Goat) and Anti-Rabbit (1:2000; Cell Signaling Technology, #7074; host: Goat). Quantification was performed by densitometry using NIH ImageJ software.

### Metabolite Extraction

For *in vivo* samples, approximately 30 mg of flash-frozen liver tissue was pulverized in 1 mL of -80°C 80:20 MeOH:H_2_O extraction solvent using a motorized tissue grinder and stored on dry ice for 5 minutes. Samples were then centrifuged at 21,100xg at 4°C for 5 minutes. Supernatants were transferred to individual 15 mL conical tubes after which 0.8 ml of -20°C 40:40:20 ACN:MeOH:H_2_O extraction solvent was used to resuspend the pelleted samples using a motorized tissue grinder. Resuspended samples were stored on wet ice for 5 minutes. Samples were then centrifuged at max speed for 5 minutes at 4°C and supernatants were combined with those collected after the initial extraction. The 40:40:20 ACN:MeOH:H_2_O extraction was then repeated as previously described. Combined metabolite extracts were centrifuged at max speed for 5 minutes at 4°C to pellet any potential insoluble debris after which supernatants were transferred to new individual 15 mL conical tubes. Finally, extracts were dried using a Thermo Fisher Savant ISS110 SpeedVac and resuspended in 150 µL H_2_O per 5 mg of the original liver input. Resuspended extracts were centrifuged at max speed for 5 minutes at 4°C after which supernatants were transferred to glass vials for LC-MS analysis.

For tissue culture samples, HepG2 cells were rapidly washed with 2x with 1 ml of ice-cold PBS pH 7.4 and incubated in 1 ml of -80°C 80:20 MeOH:H_2_O extraction solvent for 15 min at -80°C. Cells were scraped and transferred to a 1.5 ml eppendorf tube, centrifuged at 21,100xg for 5 min at 4°C, after which the supernatant was transferred to a new 2 ml eppendorf tube and stored on ice. The remaining pellet was subjected to an additional extraction by vortexing the sample with 0.8 ml of -20°C 40:40:20 ACN:MeOH:H_2_O extraction solvent. This sample was centrifuged at 21,100xg for 5 min at 4°C after which the supernatant was pooled with the 80:20 MeOH:H_2_O extraction supernatant. The remaining cell pellet was used for RIPA protein extraction and BCA protein quantification to generate normalization factors for LC-MS metabolomics data analyses. Pooled supernatants were dried using a Thermo Fisher Savant ISS110 SpeedVac and resuspended in 100 µL H_2_O. Resuspended extracts were centrifuged at max speed for 5 minutes at 4°C after which supernatants were transferred to glass vials for LC-MS analysis.

### LC-MS Metabolomics

Prepared metabolite samples were injected in random order onto a Thermo Fisher Scientific Vanquish UHPLC with a Waters Acquity UPLC BEH C18 column (1.7 μm, 2.1x100mm; Waters Corp., Milford, MA, USA) and analyzed using a Thermo Fisher Q Exactive Orbitrap mass spectrometer in negative ionization mode. LC separation was performed over a 25 minute method with a 14.5 minute linear gradient of mobile phase (buffer A, 97% water with 3% methanol, 10mM tributylamine, and acetic acid-adjusted pH of 8.3) and organic phase (buffer B, 100% methanol) (0 minute, 5% B; 2.5 minute, 5% B; 17 minute, 95% B; 19.5 minute, 5% B; 20 minute, 5% B; 25 minute, 5% B, flow rate 0.2mL/min). A quantity of 10 μ of each sample was injected into the system for analysis. The ESI settings were 30/10/1 for sheath/aux/sweep gas flow rates, 2.50kV for spray voltage, 50 for S-lens RF level, 350C for capillary temperature, and 300C for auxiliary gas heater temperature. MS1 scans were operated at resolution = 70,000, scan range = 85-1250m/z, automatic gain control target = 1 x 10^6^, and 100 ms maximum IT. Raw data files were converted into mzml for metabolite identification and peak AreaTop quantification using El-MAVEN (v0.12.1-beta) (^49^). Peak AreaTop values were imported into MetaboAnalyst for statistical analysis (one factor) using default settings (^50^).

### Transcriptomic Analysis

RNA was extracted from liver with TRI Reagent (Thermo Fisher, AM9738), then treated with DNase (Invitrogen, 12185010) and purified with RNA purification kit (Ambion, 12183025). The concentration and purity of RNA was determined using a NanoDrop 2000c spectrophotometer (Thermo Fisher Scientific, Waltham, MA) and RNA was diluted to 100-400 ng/µl for sequencing. The RNA was then submitted to the University of Wisconsin-Madison Biotechnology Center Gene Expression Center & DNA Sequencing Facility, and RNA quality was assayed using an Agilent RNA NanoChip. RNA libraries were prepared using the TruSeq Stranded Total RNA Sample Preparation protocol (Illumina, San Diego, CA) with 250ng of mRNA, and cleanup was done using RNA Clean beads (lot #17225200). Reads were aligned to the mouse (*Mus musculus*) with genome-build GRCm38.p5 of accession NCBI:GCA_000001635.7[DL9] and expected counts were generated with ensembl gene IDs (^51^).

Analysis of significantly differentially expressed genes (DEGs) was completed in R version 4.3.0. (Team, 2017) using *edgeR* (^52^) and *limma* (^53^). Gene names were converted to gene symbol and Entrez ID formats using the *mygene* package. Genes with too many missing values were removed, if genes were present in less than one diet/age group they were removed. To reduce the impact of external factors not of biological interest that may affect expression, data was normalized to ensure the expression distributions of each sample are within a similar range. We normalized using the trimmed mean of M-values (TMM), which scales to library size. Heteroscedasticity was accounted for using the voom function, DEGs were identified using an empirical Bayes moderated linear model, and log coefficients and Benjamini-Hochberg (BH) adjusted p-values were generated for each comparison of interest (^54^) (Benjamini and Hochberg, 1995). DEGs specific to each condition were rank sorted by log2 fold-change and subsequently used in GSEA Preranked analyses with default settings for gene ontology biological process (GO-BP) pathway enrichment (^34^).

### Histone Extraction and Chemical Derivatization

For *in vivo* samples, approximately 30 mg of liver tissue was resuspended in 800 μl of ice-cold Buffer A (10 mM Tris-HCl pH 7.4, 10 mM NaCl, and 3 mM MgCl2) supplied with protease and histone deacetylase inhibitors (10 μg/ml leupeptin, 10 μg/ml aprotinin, 100 μM phenylmethylsulfonyl fluoride, 10 mM nicotinamide, 1 mM sodium-butyrate, and 4 μM trichostatin A) followed by 20 strokes of loose- and 20 strokes of tight-pestle homogenization in a 1 mL Wheaton dounce homogenizer and strained through a 100 μM filter before being transferred to a new 1.5 mL Eppendorf tube. Samples were then centrifuged at 800xg for 10 minutes at 4°C to pellet nuclei. Supernatants were transferred to fresh 1.5 mL Eppendorf tubes or discarded. The remaining nuclei pellet was resuspended in 500 μl ice-cold PBS pH 7.4 and centrifuged at 800xg for 10 minutes at 4°C. The supernatant was discarded and nuclei were again washed with 500 μl ice-cold PBS pH 7.4. Next, pelleted nuclei were resuspended in 500 μl of 0.4N H_2_SO_4_ and rotated at 4°C for 4 hours. Samples were centrifuged at 3,400xg for 5 minutes at 4°C to pellet nuclear debris and precipitated non-histone proteins. Supernatants were transferred to new 1.5 mL Eppendorf tubes after which 125 μl of 100% trichloroacetic acid was added and incubated overnight on ice at 4°C. The next day, samples were centrifuged at 3,400xg for 5 minutes at 4°C to pellet precipitated histone proteins. Supernatants were discarded and precipitant was washed with 1 mL ice-cold acetone +0.1% HCl. Samples were centrifuged at 3,400xg for 2 minutes at 4°C and supernatants were again discarded. This process was repeated with a 100% ice-cold acetone wash. Residual acetone was allowed to evaporate at room temperature for 10 minutes after which dried precipitant was resuspended in H_2_O. Finally, samples were centrifuged at 21,100xg for 5 minutes at 4°C to pellet any remaining insoluble debris and supernatants containing purified histone were transferred to new 1.5 mL Eppendorf tubes. Histone extractions from flash frozen HepG2 cell pellets were performed as described above.

To prepare purified histone samples for LC-MS/MS analysis, 5 μg of each sample were diluted with H_2_O to a final volume of 9 μl. 1 μl of 1 M triethylammonium bicarbonate was added to each sample to buffer the solution to a final pH of 7-9. Next, 1 μl of 1:100 d6-acetic anhydride:H2O was added to each sample followed by a 2-minute room temperature incubation. The reaction was quenched via addition of 1 μl 80 mM hydroxylamine followed by a 20-minute room temperature incubation. Next, d3-acetylated histones were digested with 0.1 μg trypsin for 4 hours at 37°C. Upon completion of trypsin digestion, 5 μl of 0.02 M NaOH was added to adjust the final pH to ∼9-10. Prepared histone peptides were then N-terminally modified with 1 μl 1:50 phenyl isocyanate:ACN for 1-hour at 37°C. Modified peptides were desalted and eluted-off of EmporeC18 extraction membrane. Eluted samples were dried completely using a Thermo Fisher Scientific Savant ISS110 SpeedVac and resuspended in 40 μl sample diluent (94.9% H2O, 5% ACN, 0.1% TFA). Resuspended samples were centrifuged at max speed for 5 min at 4°C after which supernatants were transferred to glass vials for LC-MS/MS analysis.

### LC-MS/MS Histone Proteomics

Derivatized histone peptides were injected onto a Dionex Ultimate3000 nanoflow HPLC with a Waters nanoEase UPLC C18 column (100 m x 150 mm, 3 μm) coupled to a Thermo Fisher Q-Exactive mass spectrometer at 700 nL/ min. Aqueous phase (A) consisted of H2O + 0.1% formic acid while the mobile phase (B) consisted of acetonitrile + 0.1% formic acid (B). Histone peptides were resolved using the following linear gradient: 0 min, 2.0% B; 5 min, 2.0% B; 65 min, 35% B; 67 min, 95% B; 77 min, 95% B; 79 min, 2.0% B; 89 min, 2.0% B. Data was acquired using data-independent acquisition (DIA) mode. The mass spectrometer was operated with a MS1 scan at resolution = 35,000, automatic gain control target = 1 × 106, and scan range = 390-910 m/z, followed by a DIA scan with a loop count of 10. DIA settings were as follows: window size = 10 m/z, resolution = 17,500, automatic gain control target = 1 × 106, DIA maximum fill time = AUTO, and normalized collision energy = 30.

DIA Thermo .raw files were analyzed via the EpiProfile 2.0 AcD3 module (^55^). Subsequent data filtering (i.e., removing samples with (1) >2 null values for common peptides or (2) >50% CV) and normalization was performed using our published Histone Analysis Workflow (^56^).

### Multi-Omics Factor Analysis

To integrate transcriptomics, metabolomics, histone proteomics, and physiological data, Multi- Omics Factor Analysis (MOFA) (^35^) was performed on 60 mice consisting of 13929 transcripts, 103 metabolites, 92 histone proteins, and 15 phenotypes. The model was trained on 15 factors (minimum explained variance 0.8% in at least one data type) with default parameters except for likelihoods = “gaussian”, maxiter = 1000, convergence_mode = "slow", and seed = 42. The variance decompositions between factors and datasets were computed. Pearson correlation coefficients (Freedman et al., 2007) were calculated to show the association factors and five experimental conditions (control, isoleucine restriction, leucine restriction, methionine restriction, and rapamycin). To run an enrichment analysis, principal component gene set enrichment (PCGSE) (Frost et al., 2015) was performed on MSigDB_v6.0_C5_mouse (^57^) gene set consisting of genes annotated by GO Biological Process terms. All analyses were conducted using R (version 4.2.1) and MOFA2 (version 1.8.0).

### Statistics

Most statistical analyses were conducted using Prism, version 9 (GraphPad Software Inc., San Diego, CA, USA), R (version 4.1.0), or MatLab (version R2023b). Tests involving multiple factors were or by one-way ANOVA with Diet as the categorical variable followed by a Tukey–Kramer or Dunnett’s *post hoc* test for multiple comparisons as appropriate. Data distribution was assumed to be normal but was not formally tested.

## Acknowledgements

The Lamming lab is supported in part by the NIA (AG056771, AG081482, AG084156, and AG085898), the NIDDK (DK125859), and startup funds from UW-Madison. The Denu lab is supported in part by the NIH (GM149279 and DK125859). The Ong lab is supported in part by the NIH (U19AI104317 and DK125859), and startup funds from UW-Madison. This research was supported by The New York Stem Cell Foundation, SAH is a NYSCF - Druckenmiller Fellow. CLG was supported in part by Dalio Philanthropies, a Glenn Foundation Postdoctoral Fellowship, and by Hevolution Foundation award HF-AGE AGE-009. RB is supported by F31AG081115. The authors used the UW-Madison Biotechnology Center Gene Expression Center (RRID:SCR_017757) which is supported in part by the UWCCC (P30CA014520). The CLAMS-HC was purchased with funds from the U.S. Department of Veterans Affairs (IS1-BX005524), and this work was supported using facilities and resources from the William S. Middleton Memorial Veterans Hospital. The content is solely the responsibility of the authors and does not necessarily represent the official views of the NIH. This work does not represent the views of the Department of Veterans Affairs or the United States Government.

---

**Extended Data Figure 1:**
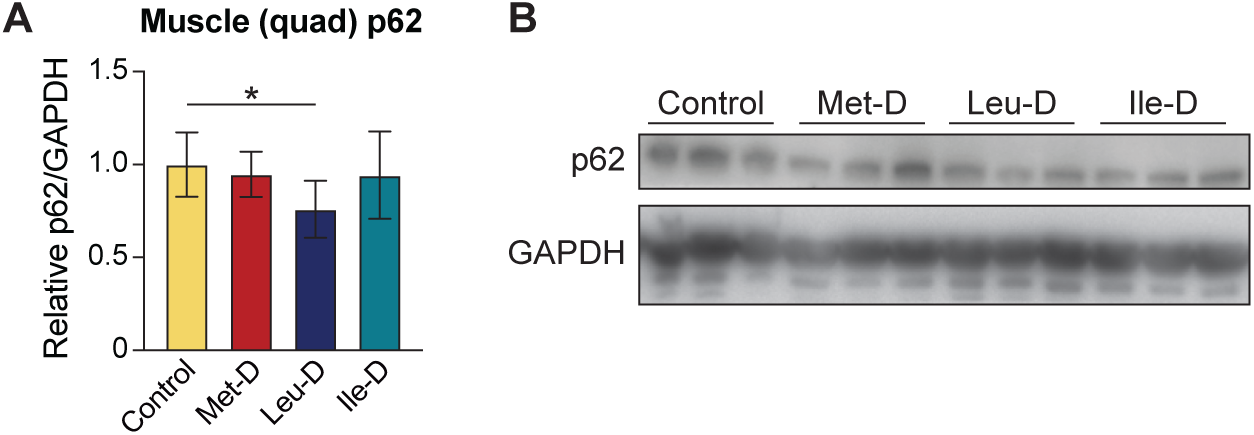
Differential p62 expression following dietary EAA depletions. (*A*) Bar graph depicting muscle (quadricep) p62 protein levels normalized to GAPDH. (*B*) Representative western blot image of those used for the quantification depicted in panel A. Error bars = SEM; N = 12; * = p-value < 0.05 as measured via Student’s t-test.

**Extended Data Figure 2: *Interactive Multi-Omics Factor Analysis Results.*** HTML file linking to interactive MOFA results.

